# Motif elucidation in ChIP-seq datasets with a knockout control

**DOI:** 10.1101/721720

**Authors:** Danielle Denisko, Coby Viner, Michael M. Hoffman

## Abstract

Chromatin immunoprecipitation-sequencing (ChIP-seq) is widely used to find transcription factor binding sites, but suffers from various sources of noise. Knocking out the target factor mitigates noise by acting as a negative control. Paired wild-type and knockout experiments can generate improved motifs but require optimal differential analysis. We introduce peaKO—a computational method to automatically optimize motif analyses with knockout controls, which we compare to two other methods. PeaKO often improves elucidation of the target factor and highlights the benefits of knockout controls, which far outperform input controls. It is freely available at https://peako.hoffmanlab.org.

## Introduction

Transcription factors, often recognizing specific patterns of DNA sequences called motifs, control gene expression by binding to *cis*-regulatory DNA elements^56^. Accurate identification of transcription factor binding sites remains a challenge^25^, with experimental noise further compounding a difficult problem^33^. Improving motif models to better capture transcription factor binding affinities at each position of the binding site facilitates downstream analyses on gene-regulatory effects. Higher-quality motifs also promote the exclusion of spurious motifs, obviating costly experimental follow-up.

Chromatin immunoprecipitation-sequencing (ChIP-seq)^30,61^ is a standard approach to locating DNA-binding protein and histone modification occupancy across the genome. Many steps of the ChIP-seq protocol can introduce noise, masking true biological signal and impeding downstream interpretation^16, 28, 33, 43, 58^. Poor antibody quality presents a major source of noise, characterized by low specificity to the target transcription factor or non-specific cross-reactivity. Cross-reactive antibodies often cause spurious pull-down of closely related transcription factor family members. Antibody clonality also contributes to antibody quality. Polyclonal antibodies tend to recognize multiple epitopes, which allows for more flexibility in binding to the desired transcription factor but at the cost of increasing background noise^33^.

To address issues of antibody quality, large consortia such as the Encyclopedia of DNA Elements (ENCODE) Project have established guidelines for validating antibodies through rigorous assessment of sensitivity and specificity^22,43^. Other considerable sources of technical noise include increased susceptibility to fragmentation in open chromatin regions^4^, and variations in sequencing efficiency of DNA segments arising from differences in base composition^33^. Downstream computational processing further reveals a different type of noise arising from contamination of peaks with zingers, motifs for non-targeted transcription factors^76^.

Additional control experiments can mitigate the effects of the aforementioned biases. Common types of controls include input and mock immunoprecipitation. Input control experiments isolate cross-linked and fragmented DNA without adding an antibody for pull down. Mock immunoprecipitation control experiments use a non-specific antibody, commonly immunoglobulin G (IgG)^28,43^, during the affinity purification step, instead of an antibody to the transcription factor. In theory, IgG mock experiments should better address technical noise since they more closely mimic the steps of the wild type (WT) ChIP protocol^43^. In practice, however, they suffer from a range of issues stemming from low yield of precipitated DNA^33^. Although the ENCODE Project22 recommends the use of input controls, these experiments also suffer from limitations. Input can only capture biases in chromatin fragmentation and sequencing efficiencies, thus failing to capture the full extent of ChIP-seq technical noise.

Knockout (KO) control experiments present an attractive alternative to input and mock immunoprecipitation. In these experiments, mutations directed to the gene encoding the target transcription factor result in little to no expression of the transcription factor, prior to ChIP-seq. This preserves most steps of the ChIP protocol, including antibody affinity purification. Therefore, KO experiments can account for both antibody-related noise and biases in library preparation.

Common transcription factor KO constructs include CRISPR/Cas9-targeted mutations17 and Cre/loxP conditional systems^66,69^. In downstream computational analyses, signal from the KO experiment serves as a negative set for subtraction from the WT positive set. Many pre-existing computational methods can use negative sets, typically input controls, to model background distributions^59, 72, 79^. For example, some peak calling tools, such as MACS2^78^, can perform discriminative peak calling. Most of these tools use the control set to set parameters of a background Poisson or negative binomial distribution5 serving as a null for assessing the significance of WT peaks^59^.

Since KO controls better account for biases inWT data than input controls, optimizing methods for KO controls should improve the quality of results from downstream analyses. Indeed, as KO constructs become increasingly more accessible^19^, the need for optimal KO processing guidelines becomes more crucial. While some preliminary studies have investigated the use of KO controls^35,49^, further rigorous comparison of methods and establishment of a standard remain necessary.

To elucidate motifs when KO controls are available, we introduce a new method, peaKO. PeaKO combines two pipelines incorporating differential processing of WT and KO datasets at different stages. By comparing the rankings of a variety of known and *de novo* motifs, we highlight peaKO’s value for discovering and assessing binding motifs of WT/KO experiments, and peaKO’s applications in other differential contexts.

## Results

### PeaKO combines two differential analysis pipelines

Two steps of ChIP-seq computational processing allow for the subtraction of control signal from WT signal: peak calling and motif analysis. Therefore, we created two complementary pipelines, Pipeline A and Pipeline B, integrating the same software tools but selecting opposing steps to subtract matched KO signal from WT signal. (Figure 1A).

**Figure 1.**
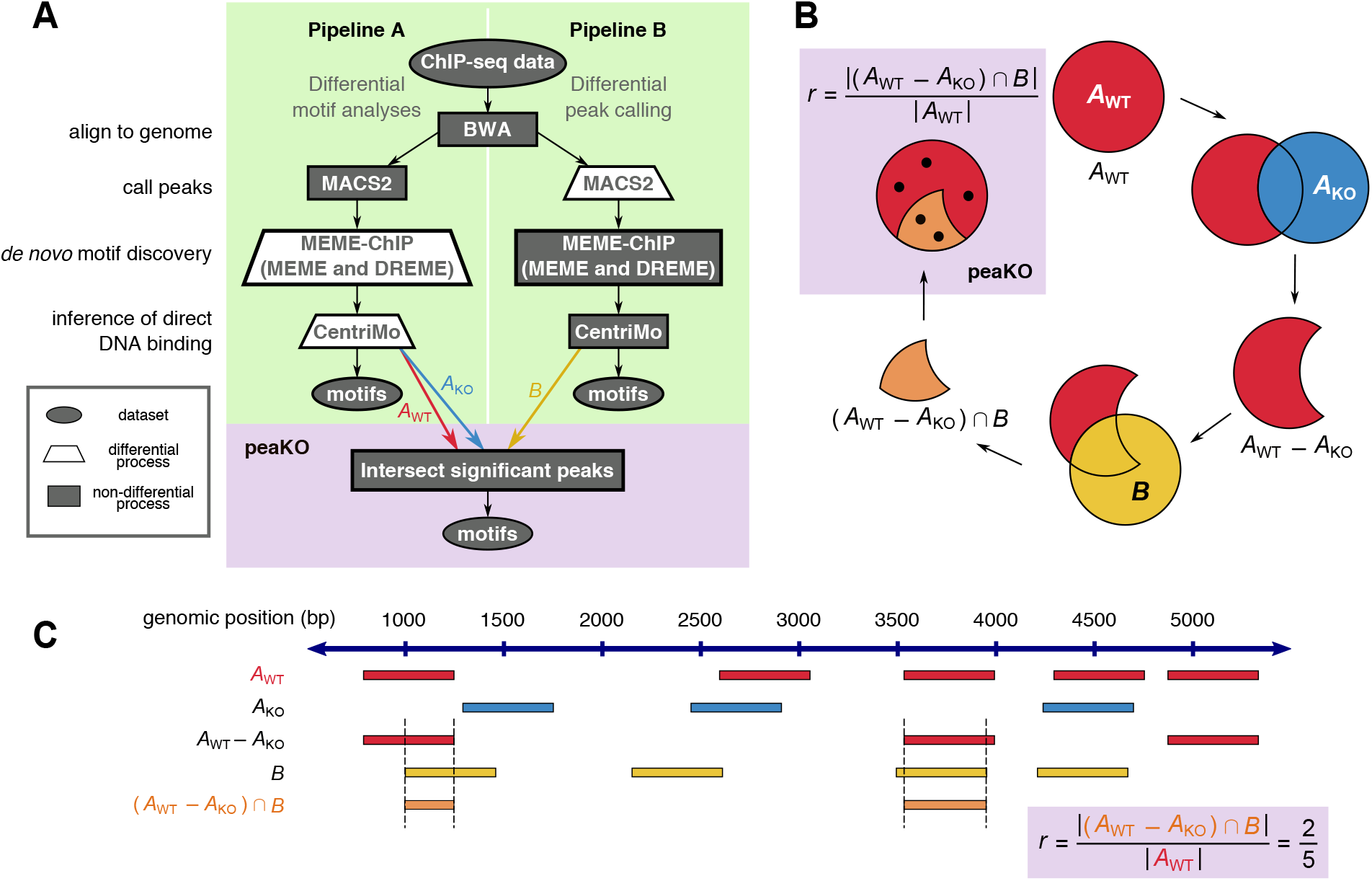
Overview of Pipelines A and B, and peaKO. **(A)** Pipelines A and B differ in their differential analysis steps. Each pipeline accepts both wild type (WT) and knockout (KO) ChIP-seq data as input. Pipeline A incorporates differential motif elucidation via MEME-ChIP^51^, whereas Pipeline B incorporates differential peak calling via MACS2^78^. Both pipelines produce a ranked list of motifs predicted as relevant to the ChIP-seq experiment by CentriMo^9,44^. PeaKO extracts significant peaks from CentriMo and computes a new score by which it ranks motifs. **(B)** PeaKO computes its ranking metric *r* through a series of set operations. PeaKO uses peak sets *A*_WT_ and *A*_KO_, extracted from Pipeline *A*, and peak set *B,* extracted from Pipeline B. **(C)** A toy example illustrates the calculation of peaKO’s score. Starting from the top row of peak set *A*_WT_ and moving downwards, we apply the peak set operations of *r* sequentially to identify regions satisfying the numerator criteria.

Pipeline A incorporates differential motif analysis through MEME-ChIP^50,51^. It focuses on the motif discovery algorithms MEME^7,8^ and DREME^6^, and includes the motif enrichment algorithm CentriMo^9,44^. MEME-ChIP uses control peak sets for discriminative enrichment analysis^50^.

Instead of differential motif analyses, Pipeline B incorporates differential peak calling through MACS2^78^. MACS2 uses the control peak set to set the parameters of the background null distribution from which it calls significant peaks. Pipeline B drew inspiration from the knockout implemented normalization (KOIN) pipeline^35^.

Both pipelines conclude by executing CentriMo^9,44^. CentriMo’s measure of motif central enrichment assesses the direct DNA binding of the enriched transcription factor^9^. Some aspects of CentriMo’s output differ according to whether we choose differential44 or non-differential9 mode. Both pipelines, however, output a list of motifs ranked in order of increasing p-values. Ideally, the top motif should reflect the target factor in the underlying ChIP-seq experiment, although some circumstances may preclude this.

Each pipeline incorporates a unique approach to discriminative analysis. By modeling the peak background distribution using the negative control set, Pipeline B directly compares the position of read pileups between positive and negative datasets. In this model, we assume that read pileups shared between both datasets represent technical noise, while the remaining significant WT read pileups represent binding of the target transcription factor. Conversely, Pipeline A disregards the positional information of peaks and instead focuses on the position of the motif matches within the peaks. Pipeline A takes into account each peak’s membership in the positive or negative set only when assessing the statistical significance of a motif. In Pipeline A, the simple motif discovery tool DREME compares the fraction of *de novo* motif matches in WT sequences to KO sequences. We assume that motifs more often located near peak centers in the WT dataset than in the KO dataset suggest associated binding events.

To select for motifs that both have consistent matches within peaks and fall within regions of significant read pileup, we combined both pipelines in a new way to develop peaKO. For each motif, peaKO computes the number of overlapping peaks between peak sets generated by both pipelines, with overlaps interpreted as genuine binding events (Figure 1B and Figure 1C; see Methods).

### PeaKO usually improves or maintains the best ranking of the known motif

To assess the performance of each method, we can first compare how well methods rank known canonical motifs of sequence-specific transcription factor datasets. We collected publicly available WT/KO paired ChIP-seq datasets for 8 sequence-specific transcription factors: ATF3^80^, ATF4^27^, CHOP^27^, GATA3^74^, MEF2D^3^, OCT4^34^, SRF^70^, and TEAD431 (Table 1). We evaluated our methods on these datasets, supplementing CentriMo with the collection of vertebrate motifs from the JASPAR 2016 database53 (see Methods). Each transcription factor in our WT/KO datasets contains a corresponding motif within the JASPAR database. We used these JASPAR motifs as our gold-standard known motifs, and compared their rankings across methods. As a control, we processed theWT dataset alone through the same pipeline steps without any KO data.

**Table 1.**
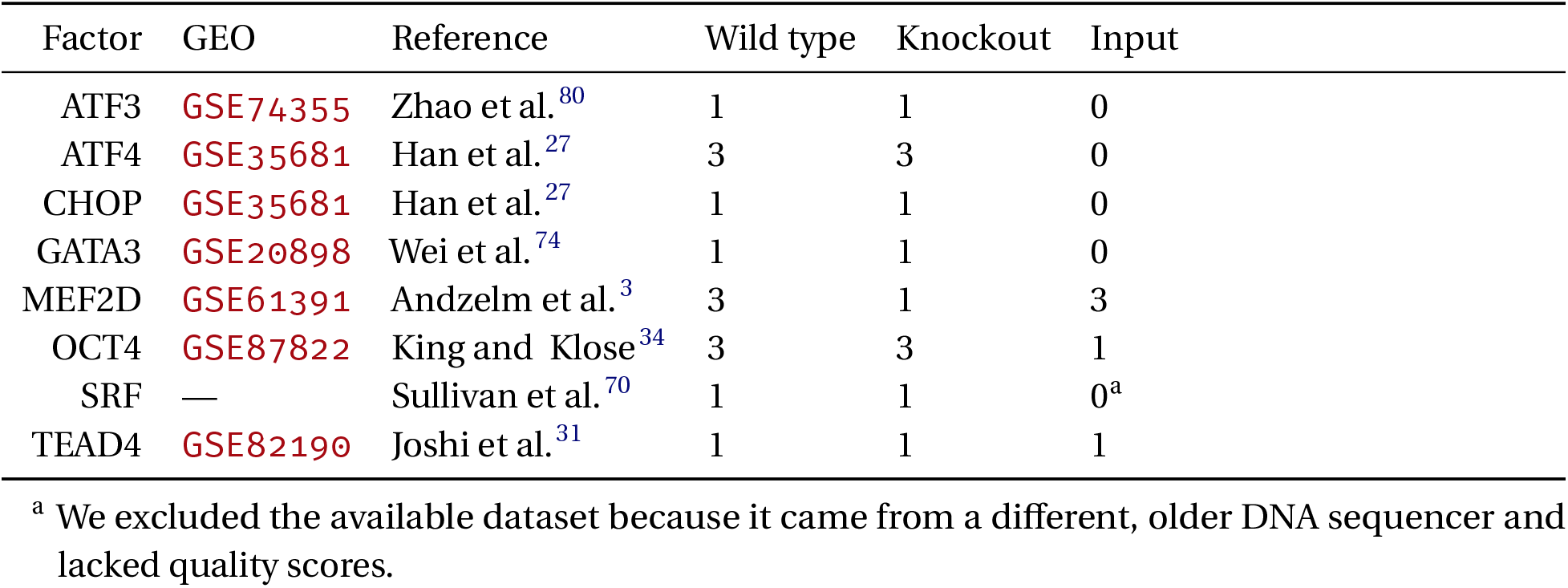
ChIP-seq datasets used, with associated Gene Expression Omnibus (GEO) accession numbers (where applicable) and number of replicates.

In 5 out of 8 cases, peaKO improved or maintained the optimal rank relative to all other methods. PeaKO also always improved or maintained the rank relative to at least one other method (Figure 2). The total number of ranked motifs differed between experiments, which suggests peaKO may benefit analyses for a wide range of transcription factors with variable binding affinities. Of the other methods, Pipeline A performed the worst overall, as exemplified by non-significant Fisher E-values for both the GATA3 and ATF3 datasets. Pipeline B performed similarly to the use of only WT data processed without controls, suggesting it benefits little from the control. PeaKO combines the best aspects of both types of differential analysis pipelines, limiting their deficiencies and highlighting their strengths. This generally leads to better rankings of known motifs.

**Figure 2.**
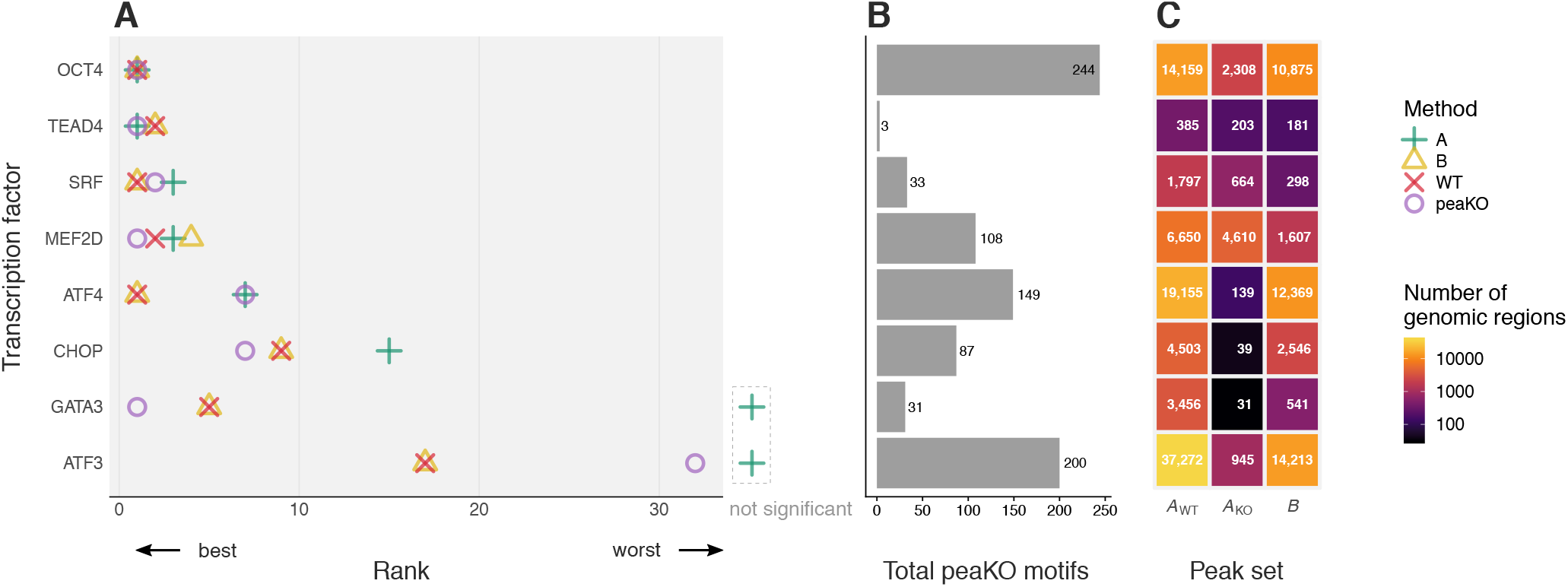
Known transcription factor motifs elucidated by different methods. Motifs originated from the JASPAR 2016 motif database^53^. Knockout datasets served as a control for differential analyses. **(A)** Each method ranked JASPAR database motifs based on their centrality within peak sets, as determined by CentriMo^9,44^. Ranks correspond to the ChIPped transcription factor’s known motif (Table 2). Dashed area to right of plot: motifs without statistical significance. **(B)** Total number of motifs assessed by peaKO. **(C)** The number of peaks found by each method varies across peak sets.

### *De novo* motifs consistently match known motifs

We investigated each method’s ability to rank *de novo* motifs and assessed the similarity between *de novo* and known JASPAR motifs. For consistency, we pooled *de novo* motifs generated by each method (see Methods). We quantified similarity between *de novo* and known motifs using Tomtom^26^. We studied these methods on the same 8 WT/KO paired datasets used for our known motif analyses.

Usually, top *de novo* motifs more closely resembled the canonical motif across methods, resulting in most ranking near 1 (Figure 3). Conversely, motifs ranking lower tended to have fewer matches to the known motif, often not even matching the known motif at all. PeaKO generally followed this trend, but in a few exceptions, such as CHOP, OCT4, and ATF3, top motifs also sparsely matched the canonical motif. PeaKO might have found related, interacting factors, rather than the factor of interest. For example, many top *de novo* motifs reported by peaKO for the CHOP dataset closely matched the motif for ATF4, which interacts with CHOP^27^.

**Figure 3.**
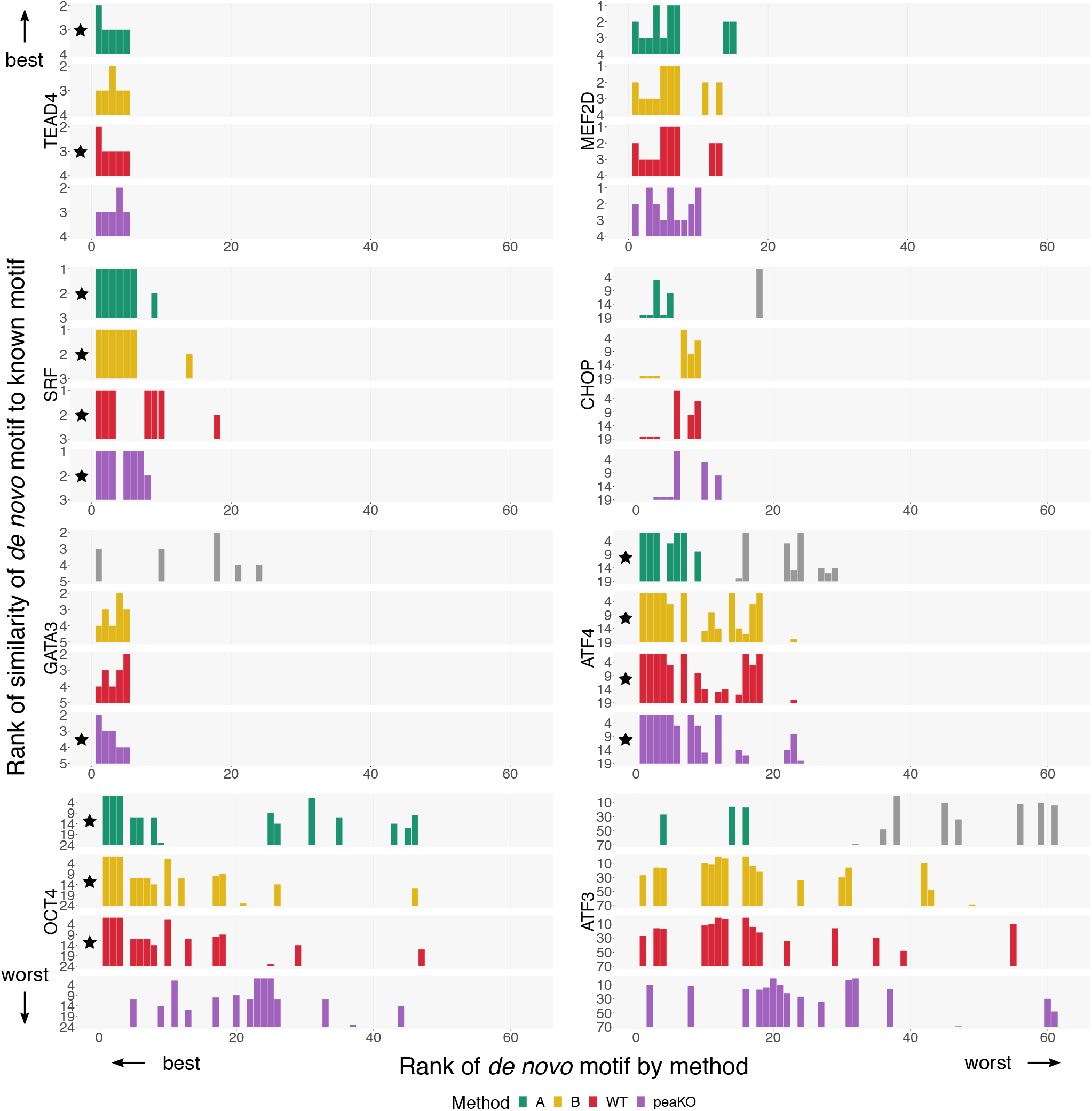
Similarity of discovered *de novo* motifs to canonical JASPAR motifs across 4 methods. For 8 transcription factors (Table 2), we ran 4 methods (green: Pipeline A, yellow: Pipeline B, red: WT alone, purple: peaKO) on a pooled set of *de novo* motifs generated by MEME^7,8^ and DREME^6^. Each method generated a ranking of *de novo* motifs (x-axis). For each of these motifs, we quantified similarity to the known motif using Tomtom^26^ (y-axis). For a given *de novo* motif, Tomtom finds and ranks the most similar motifs from the total set of JASPAR motifs. We record the rank of the transcription factor’s known JASPAR motif within this list. To emphasize strong matches to known motifs, the provided ranks lie in descending order, with the best (rank 1) motif, at the top. In some cases, the best rank achieved by the match does not reach 1, as reflected by a greater lower limit. Black stars: methods achieving the best possible rank (of 1, for method rankings and the lower limit rank, for similarity rankings) across both ranking schemes within each experiment. Gray bars: motifs that were not statistically significant and gave rise to arbitrary rankings.

### PeaKO teases apart similar GATA family motifs

We delved deeper into our GATA3 results, for which peaKO outperformed all other methods. GATA3 belongs to the family of GATA factors, all of which bind GATA-containing sequences^55^. Despite having similar motifs, each GATA factor plays a distinct role and usually does not interact with the others^73^.

Distinguishing the targeted motif among GATA factors and other large transcription factor families often presents a challenge. Minor differences in position weight matrices (PWMs)^14^ can cause major differences in genome-wide transcription factor binding sites^40^. Understanding the downstream effects of transcription factor binding necessitates pulling apart these intricacies in motif preferences.

CentriMo results across both pipelines further reinforced the difficulty of distinguishing these motifs (Figure 4). Pipeline B identified closely related GATA family members with ranks 1–4, above the desired fifth-ranked GATA3 motif. Pipeline A proved less promising, failing to reliably rank any GATA family members. Furthermore, none of the shown Pipeline A motifs appeared centrally enriched within WT peaks. Instead, we observed a uniform distribution among the WT peak set and a series of stochastic, sharp peaks among the KO peak set, likely representing inflated probabilities due to low sample size.

**Figure 4.**
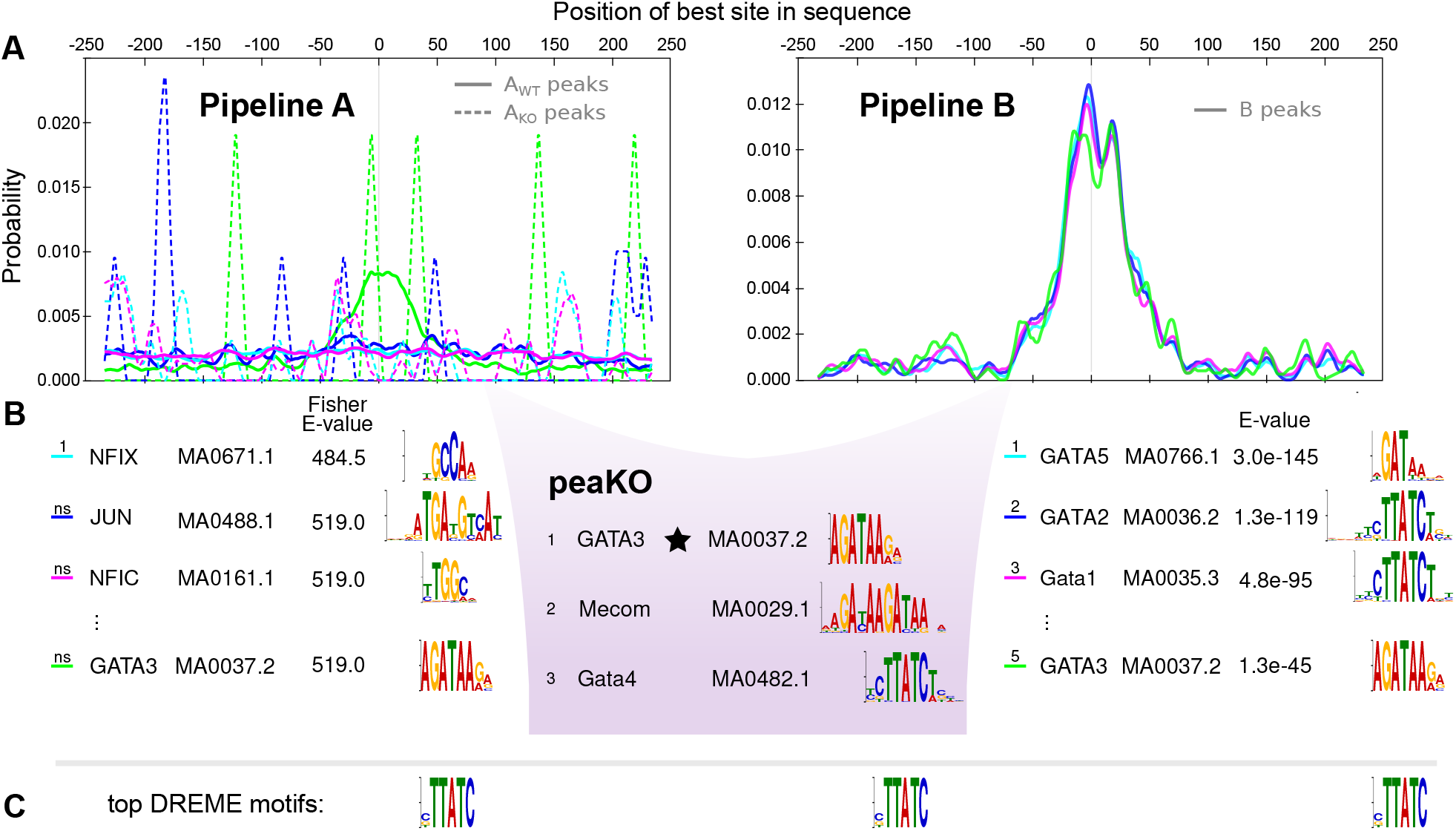
PeaKO ranks the GATA3 motif above other GATA transcription factor family motifs. **(A)** CentriMo^9,44^ probability plots depict enrichment of the top 3 motifs from each method, along with the GATA3 motif, within peak sets. **(B)** Motifs resulting from each pipeline and peaKO lie beneath associated CentriMo plots. Motifs and corresponding sequence logos65 originate from JASPAR 2016^53^. Capitalization is as it occurs in JASPAR. The information content of bases in the sequence logos ranges from 0.0000 bits to 2.0000 bits. Ranks of “ns” indicate non-significant motifs, and therefore unreliable rankings. The black star denotes achieving the best rank of the GATA3 motif. **(C)** Top DREME6 motifs with length greater than 5.0000 bp, for comparison. In this case, all three motifs are identical.

Despite the difficulties affecting Pipeline B, peaKO draws on its ability to detect GATA family members, and surpasses it by ranking GATA3 first. In this case, peaKO achieves specificity in ranking motifs in the presence of many similar familial motifs.

### Low-quality datasets account for poor rankings across methods

In a few cases, peaKO performed worse than the other methods at ranking the canonical motif (Figure 2). In particular, we observed a large spread in rankings across methods for ATF3 (ranging from rank 17 to non-significant ranks, above 80). We found central enrichment of the canonical ATF3 motif in the KO peak set, as depicted by Pipeline A’s CentriMo results (Figure 5). This central enrichment appears even more prominent than that in the WT peak set.

**Figure 5.**
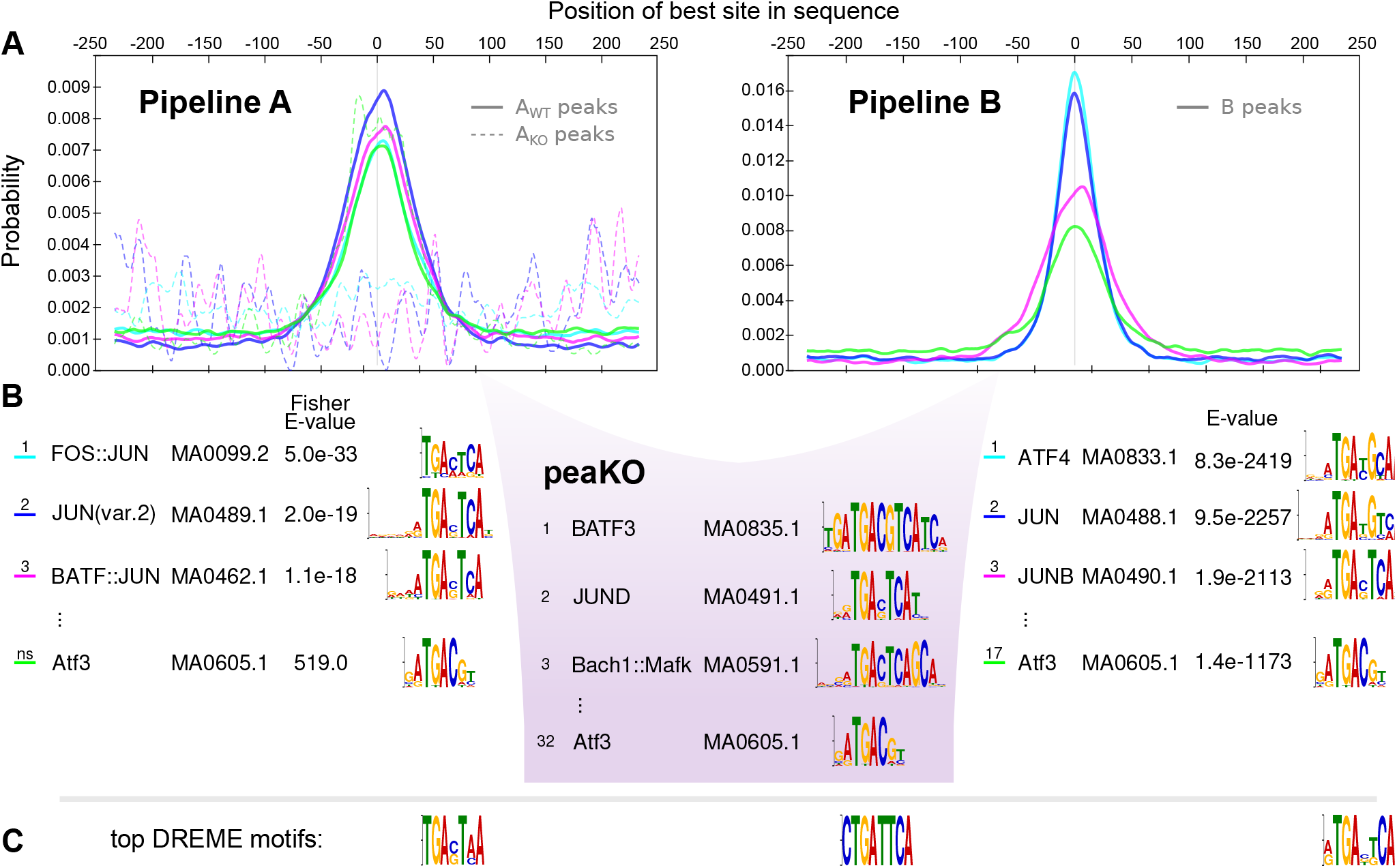
The ATF3 motif is centrally enriched in the ATF3 knockout dataset. **(A)** CentriMo^9,44^ probability plots depict enrichment of the top 3 motifs from each method, along with the ATF3 motif, within peak sets. **(B)** Motifs resulting from each pipeline and peaKO lie beneath associated CentriMo plots. Motifs and corresponding sequence logos65 originate from JASPAR 2016^53^. Capitalization is as it occurs in JASPAR. Information content of bases underlying motifs range from 0.0000 bits to 2.0000 bits. Ranks of “ns” indicate non-significant motifs, and therefore unreliable rankings. **(C)** Top DREME6 motifs with length greater than 5.0000 bp, for comparison.

Although CentriMo probabilities depend on the total number of peaks in each set, and a relatively low number of peaks in the control set can inflate these probabilities, we expect non-specific matches to generate a uniform background distribution rather than a distinctive centrally-enriched pattern^9,44^. Accordingly, ATF3 enrichment deviates substantially from our expectations and suggests issues with the underlying KO ChIP experiment. This likely explains the poor rankings of ATF3 across methods, including peaKO.

### Knockout-controlled analyses consistently improve motif elucidation

To investigate whether KO controls would better approximate WT ChIP-seq experimental noise than input controls, we used input controls to repeat our analyses. We ran our methods on MEF2D, OCT4, and TEAD4 datasets, which contained input controls (Table 1), by applying the same procedures but using only the input dataset for differential analysis steps.

Using an input control instead of a KO control usually worsened the ranking of the known motif, as observed by an overall shift across methods toward poorer rankings (Figure 6A). In *de novo* motif analyses with input controls, top-ranked motifs tended to have slightly poorer matches to known motifs across methods, as compared with KO controls (Figure 6B). As in WT/KO analyses of OCT4, we observed sparsity in top-ranked peaKO motifs matching the known motif. This could point to low affinity of the antibody to the target factor or other types of noise affecting primarily the WT set. Indeed, input experiments yielded even fewer significant peaks from CentriMo than KO experiments (Figure 6C).

**Figure 6.**
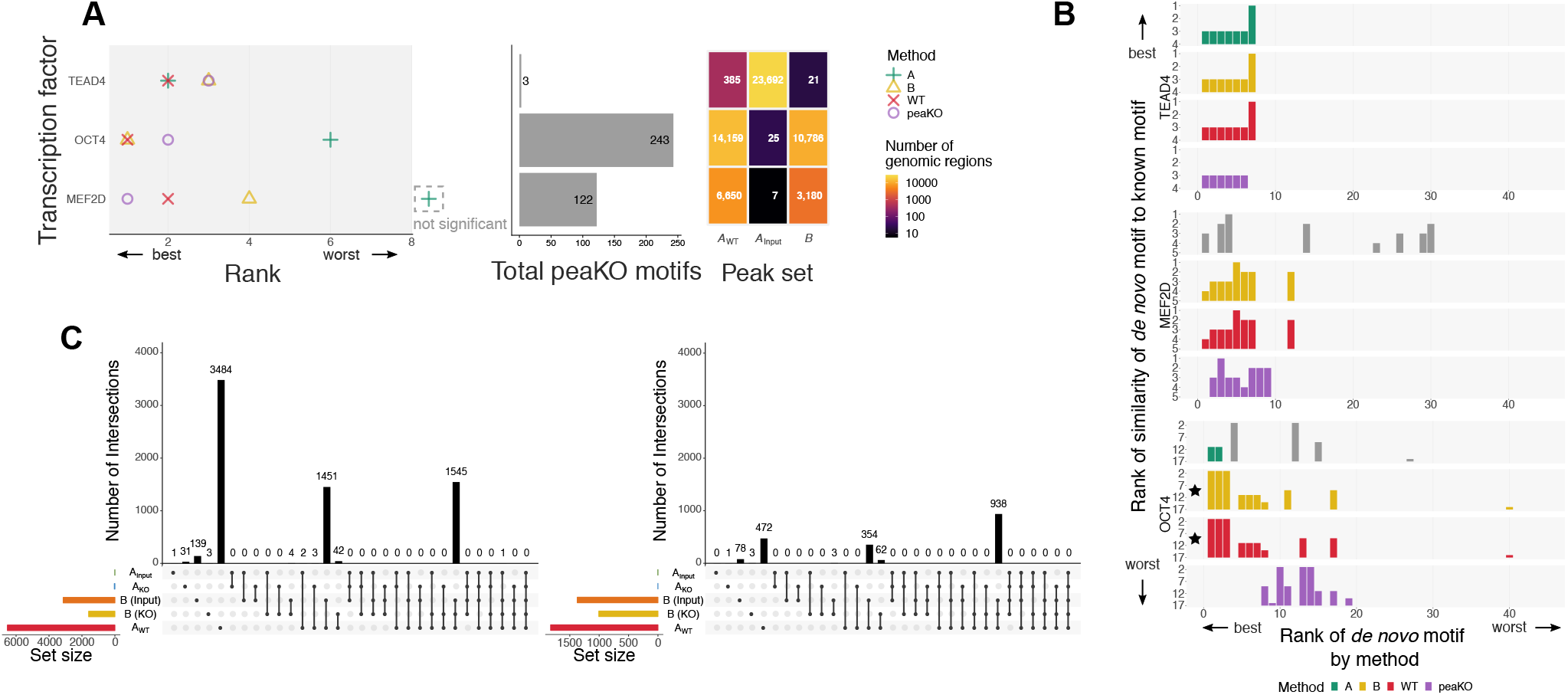
Ranks of known and discovered motifs using input controls. **(A)** Ranks of known JASPAR53 motifs across methods for each ChIP-seq experiment (Table 2). Input datasets served as a control in differential analysis steps. Dashed area to right of plot: motif without statistical significance. **(B)** We plotted ranks of *de novo* motifs discovered by MEME^7,8^ and DREME6 against their similarity to the known JASPAR motif, as quantified by Tomtom^26^. We compared queried motifs against the JASPAR 2016 target motif database. Gray bars: motifs that were not statistically significant and gave rise to arbitrary rankings. Black stars: methods achieving the best possible rank (rank of 1 for method rankings and lower limit rank for similarity rankings) across both ranking schemes within each experiment. **(C)** UpSet plot45 of overlap between MEF2D peak sets generated by Intervene^32^ *(left*) for all motifs and (*right*) for the MEF2D motif only. For Pipeline B peak sets, parentheses indicate the type of negative control used for peak calling: input or knockout (KO).

Overall, using input controls instead of KO controls led to poorer rankings across methods. Although peaKO did not outperform the other methods using only input, it generally performed similarly, suggesting utility in other differential applications.

## Discussion

Increased accessibility of KO experiments presents a need for standardized computational processing workflows. With KO data, peaKO’s dual pipeline approach generally outperformed each pipeline alone when ranking the known motifs. This holds true even in challenging cases, such as distinguishing among large transcription factor families with shared core motifs. Applying our methods to datasets containing both input and KO controls demonstrates the superiority of KO controls for motif elucidation.

We observed a common theme throughout our analyses pertaining to the characteristic performance of each pipeline alone. When tasked with ranking the known motif, Pipeline A generally produced inferior rankings, especially for ATF3 and GATA3 (non-significant ranks of canonical motifs) and, to a lesser extent, CHOP (rank 15). We could only attribute this to poor experimental quality for ATF3. The significance of differential mode CentriMo p-values, calculated using Fisher’s exact test, appears closely linked to the relative size of each peak set. Both CHOP and GATA3 KO control sets had fewer than 50 KO peaks (Figure 2), which might account for Pipeline A’s poor performance.

Pipeline B suffered from a different issue: it ranked known motifs almost identically to WT processing alone, without any controls. In some cases, WT processing alone even surpassed Pipeline B. WT-only processing out-performed Pipeline B when using KO and input controls for MEF2D, and when using input controls for TEAD4. Since the sole difference between Pipeline B and WT-only processing lies in the peak calling step, identical rankings indicate the sufficiency of constructing the background distribution with WT-derived values alone. Indeed, similar rankings may point to a shortcoming in comprehensively modeling noise captured in KO and input controls. Future work should investigate the robustness of peak callers in modeling control signals and explore potential integration of other tools designed for identifying differential peak sets, such as DiffBind^68^. Differential peak calling with KO controls does, however, reduce the size of theWT peak set. Perhaps this improves an already specific peak set such that the improvement is largely undetectable when ranking known motifs. Nonetheless, rankings differ in some cases and *de novo* motif analyses reveal differences between Pipeline B and WT-only processing. Overall, both pipelines show strengths in specific contexts, which peaKO emphasizes.

In selecting known canonical motifs as ground truths to assess our experiments, we limited our ability to evaluate each method’s detection of higher quality motifs. We partially addressed this limitation by finding *de novo* motifs in a discovery use case. Maintaining consistency in performance evaluation across analyses, however, required comparisons of *de novo* motifs to their cognate JASPAR motifs. Therefore, even in the discovery use case, we remain limited by the quality of the known canonical motifs.

Some of our methods ranked the motif of interest less favorably than other GATA family member motifs. GATA family members share a common core motif, yet each have distinct and detectable binding preferences that contribute to their diversity in genome-wide occupancy and function^55^. Finding the general familial motif could prove sufficient in some cases^63^. Nonetheless, finding the specific motif helps with understanding the roles of individual transcription factors.

The GATA3 motif (MA0037.2)^53^ that we ranked first (Figure 4) was generated from 4628 curated sites from a ChIP-seq experiment. This motif likely has more similarity to actual GATA3 binding sites than the other GATA family motifs we compared against, in that it was elucidated via an antibody attempting to specifically target GATA3. CentriMo selected this particular motif as the most enriched match, choosing it over other GATA3 motifs in JASPAR. Even if this motif does not model the full intricacies of GATA3 binding, one would still prefer it ranked above motifs from assays targeting other GATA family members. Our ability to rank the target motif first mainly provides additional confirmation that peaKO performs well and likely has utility in other contexts, including those focused on differentiating similar motifs.

For OCT4 (also known as POU5F1), we selected the Pou5f1::Sox2 motif (MA0142.1). SOX2, like OCT4, regulates pluripotency in embryonic stem cells^77^. The two transcription factors often act together to regulate gene expression by forming a complex and co-binding to DNA^1^. Here, however, the heterodimer motif differs substantially from the OCT4 motif alone, as it additionally contains a SOX2 motif^1^. We chose to use the heterodimer motif in assessing our methods because the authors of the study that generated the OCT4 dataset found a substantially larger proportion of peaks containing the heterodimer (44.0%) as compared to the monomer (20.6%)^34^. Upon re-running our analyses using the monomer motif instead, we found poorer rankings across methods, as expected from this imbalance of motif types in peaks (see https://doi.org/10.5281/zenodo.3338330). Higher occupancy of the heterodimer form, however, does not preclude the transcription factor from binding DNA in its monomer form. Although all methods found the heterodimer motif as the top rank, deciding upon which motif form to use and how it affects downstream processing would benefit from further exploration.

We highlighted the ATF3 experiment as a case where peaKO performs poorly, and attributed the poor performance largely to the dataset’s poor experimental quality. Nonetheless, the low information content of the ATF3 motif used as ground truth may supply an alternative explanation. In the JASPAR 2016 motif database, ATF3 motif MA0605.1 captures a single core `TGAC` motif^53^. A newer version of this motif (MA0605.2, added in JASPAR 2020^23^), however, contains the canonical ATF/CRE motif, which appears to better represent ATF3’s typical *in vivo* homo- or heterodimerization^62^. Many of the top motifs returned by peaKO and other methods appear similar to this newer version. This suggests that peaKO may have performed better than we were able to assess with JASPAR 2016, and highlights the benefit of peaKO for motif discovery.

Our use of cross-species PWMs potentially limits our findings. We used motifs from the JASPAR vertebrate collection interchangeably where the known motif did not always originate from the same species as our ChIP-seq datasets (see Tables 1 and 2). Recently, Lambert et al.42 found that, contrary to commonly held belief, extensive motif diversification among orthologous transcription factors occurs quickly as species diverge. Additionally, PWMs14 themselves, while providing the most commonly used motif model^12,37^, may not sufficiently capture nuanced binding differences^20,37^.

**Table 2.** Known JASPAR CORE 2016^53^ vertebrate motifs. Sentence case motif names designate mouse transcription factor motifs, while full upper case names designate human motifs. Double colons designate heterodimer motifs.

| Common name | JASPAR ID | Motif name |
| --- | --- | --- |
| ATF3 | MA0605.1 | Atf3 |
| ATF4 | MA0833.1 | ATF4 |
| CHOP | MA0019.1 | Ddit3::Cebpa |
| GATA3 | MA0037.2 | GATA3 |
| MEF2D | MA0773.1 | MEF2D |
| OCT4 | MA0142.1 | Pou5f1::Sox2 |
| SRF | MA0083.3 | SRF |
| TEAD4 | MA0809.1 | TEAD4 |

Technical biases can negatively influence peaKO’s scoring metric. Antibody quality, sequencing biases, and various batch effects, can lead to a failure to recover sufficient peaks or can enrich for additional off-target peaks. This can, in turn, alter the proportion of peaks in peaKO’s score, changing the motif ranking, in a potentially confounded manner. Further work is needed to assess the statistical robustness of peaKO’s score in the face of such biases. Even without that assessment, our empirical results demonstrate that peaKO’s score remains useful. Furthermore, as we have previously discussed, ranking discrepancies between motifs become obvious in peaKO results. PeaKO, like the vast majority of software that works to elucidate transcription factor binding sites, requires sequence-specific transcription factors that are suitable for narrow peak calling. While this includes the majority of transcription factors, it implies that this method is not applicable for the analysis of histone marks or other broad peak targets. Irrespective of improved motif rankings, peaKO facilitates differential motif comparisons and the generation of potentially improved *de novo* motifs.

Lastly, we used peaKO along with our other methods to assess the benefit of KO controls over input, suggesting that peaKO may prove useful for other non-WT/KO differential contexts. CRISPR epitope tagging ChIP-seq (CETCh-seq), which involves the insertion and expression of FLAG epitope tags on the target transcription factor^64^, presents one alternative differential context which may gain from peaKO. CETCh-seq provides a substantial advantage over traditional ChIP-seq because it only requires one high-quality monoclonal antibody recognizing the FLAG antigen across any number of transcription factor experiments. Notably, CETCh-seq also avoids artefacts that may arise from perturbations of the cellular context in knockout experiments. Preliminary analyses using CETCh-seq datasets revealed challenges arising from unexpected signal from a shared control of ChIP-seq in an untagged cell line. Further work should investigate the role of CETCh-seq controls and how they integrate with peaKO. We expect this work to also prove useful for comparable or enhanced methods, like CUT&RUN^67^, which one could similarly improve through careful design of complementary controls and differential motif analyses.

Similar considerations for the proper use of control sets could also apply to combining replicates. Combining negative control replicates with the irreproducible discovery rate (IDR) framework48 may pose problems considering that these datasets represent noise rather than a full range across true signal and noise. This may present an issue as IDR’s underlying copula mixture model assumes the existence of an inflection point within the dataset marking the transition between true signal and noise^48^.

## Conclusion

We present peaKO, a free and publicly available tool for ChIP-seq motif analyses with KO controls (https://peako.hoffmanlab.org). PeaKO improves over two kinds of differential processing in ranking the motif of interest. We anticipate that peaKO will prove useful in identifying motifs of novel transcription factors with available KO controls. We hope this will encourage both greater collection and wider usage of knockout datasets.

## Methods

### Overview of ChIP-seq processing and analysis methods

ChIP-seq processing follows this overarching path:

1. subject sequenced reads to trimming and quality control assessment;
2. align reads to a reference genome;
3. call peaks according to significant read pileups;and
4. elucidate *de novo* motifs and assess peaks for evidence of direct DNA binding.

For some methods, steps 3 and 4 can incorporate information from control datasets. We constructed two pipelines to compare differential analyses in both of these steps (Figure 1A).

In Pipeline A, we perform differential analysis with MEME-ChIP^50,51^. MEME-ChIP uses the *de novo* motif elucidation tools MEME^7,8^ and DREME^6^, and assesses the central enrichment of motifs in peaks via CentriMo^9,44^. CentriMo ranks motifs according to multiple-testing corrected binomial p-values (non-differential mode)^9^ or Fisher’s exact test p-values (differential mode)^44^.

In Pipeline B, we perform differential peak calling through MACS2^78^. While Pipeline B draws inspiration from the KOIN pipeline^35^, it does not incorporate the HOMER `makeTagDirectory` or `annotatePeaks`^29^ steps. We replaced HOMER motif tools29 with those from the MEME Suite^10,11^. Both Pipelines A and B incorporate identical pre-processing and alignment steps, described later. Since both pipelines employ CentriMo in their last step, they generate a list of ranked motifs with predicted association to the ChIP-seq experiment.

### PeaKO: motivation and score

Differential peak calling and differential motif analysis address the same problem of noise removal, albeit in distinct ways. Therefore, we surmised that by combining the two approaches, the results from each pipeline could complement and strengthen one another. CentriMo produces a ranked list of motifs, and each motif has an associated peak set containing a centered window enriched for that motif. We reasoned that motifs with a large proportion of peaks shared between both pipelines are likely relevant to the ChIP-seq experiment. We then created a metric that captures this.

PeaKO takes as input the CentriMo output of each pipeline. We modified CentriMo code to output negative control set peaks associated with each motif in differential mode, since at the time of this work, the software only output positive peaks. These changes have now been incorporated into the MEME Suite, and are available from all versions since 5.0.0. From the CentriMo results peaKO filters out motifs with multiple-testing corrected p-values > 0.1.

PeaKO computes a ranking metric *r* that represents the proportion of high-quality *A*_WT_ peaks found in set *B* but not in set *A*_KO_. To do this, peaKO calculates the overlap between peak sets *A*_WT_ and *A*_KO_ from Pipeline A, and peak set *B* from Pipeline B through a series of set operations:

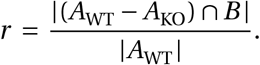

PeaKO performs this by employing pybedtools (version 0.7.7; BEDTools version 2.26.0)^18,60^. First, peaKO removes any *A*_WT_ peak overlapping at least 1.0000 bp of an *A*_KO_ peak(`pybedtools subtract-A`; Figure 1B). Second, peaKO finds regions overlapping by at least 1.0000 bp between remaining *A*_WT_ peaks and *B* peaks (`pybedtools intersect -wa`). Third, peaKO applies pybedtools merge with default settings to overlapping regions, which merges identical regions and ensures that the ranking metric *r* has a maximum value of 1. PeaKO’s final output consists of a list of motifs ranked according to this metric.

### Datasets

We analyzed a total of 8 publicly available ChIP-seq experiment datasets with KO controls (Table 1). Of these 8, we selected two datasets (GATA3 and SRF) from Krebs et al.^35^, while we selected the remainder by searching for KO-associated ChIP-seq datasets on GEO^21^. We accessed datasets through GEO, except for the SRF dataset^29^, available on Zenodo (https://doi.org/10.5281/zenodo.3405482). ATF3 experiments come from human tissue, while the other experiments come from mouse tissue.

### Motifs

We downloaded the collection of vertebrate motifs in MEME format11 from the JASPAR CORE 2016 motif database, which consists of curated PWMs derived from *in vivo* and *in vitro* methods^53^.

We defined each canonical motif from the JASPAR collection as the motif matching the target transcription factor except in two cases: OCT4 and CHOP (Table 2). In both cases, we instead chose motifs derived from their common heterodimer complex forms. CHOP or DDIT3 likely binds DNA as an obligate multimer^41,57^, so we used Ddit3::Cebpa (MA0019.1). The CHOP monomer motif closely resembles its C/EBPα heterodimer motif, relative to its Cis-BP (version 1.02)^75^ DDIT3 motif (`T025314_1.02`, derived from HOCOMOCO^38^). For OCT4, we used the Pou5f1::Sox2 motif (MA0142.1; see Discussion).

We provided motifs to CentriMo^9,44^ for central enrichment analyses and to Tomtom26 for similarity assessments.

### Pre-processing, alignment, and peak calling

Before alignment, we trimmed adapter sequences with TrimGalore! (version 0.4.1)^36^ which uses Cutadapt (version 1.8.3)^52^. We assessed sequencing data quality using FastQC (version 0.11.5)^2^. We used Picard’s `FixMateInformation` and `Add0rReplaceReadsGroups` (version 2.10.5)^15^ and GATK’s `PrintReads` (version 3.6)^54^ to prevent GATK errors. Then, we aligned reads to GRCm38/mm10 or GRCh38/hg38 with BWA `bwa-aln` (version 0.7.15)^46^ (as recommended^47^, since some datasets have reads << 70.0000 bp), using Sambamba (version 0.6.6)^71^ for post-processing.

Next, we called peaks using MACS2 (version 2.0.10)^78^ with parameters `-q 0.05`. In Pipeline A, we called WT and KO peaks separately. In Pipeline B, we provided the KO dataset as a control to the WT dataset during peak calling (parameter -c), resulting in a single set of peaks.

### Combining replicates

For MEF2D, OCT4, and TEAD4 experiments which consist of biological replicates (see Table 1), we processed replicates using the ENCODE Transcription Factor and Histone ChIP-seq processing pipeline^39^. The ENCODE pipeline replaces the pre-processing, alignment, and peak calling steps described earlier. We chose default parameters for punctate (narrow peak) binding experiments in all steps. Instead of a q-value threshold, this pipeline caps the number of peaks (*n* = 500000.0000) to ensure that the IDR framework48 can analyze a sufficient number of peaks across a full spectrum. IDR combines peaks across replicates based on the assumption that strong peaks shared across replicates represent true binding events, while weak, one-off peaks represent noise. To emulate the first steps of Pipeline A and Pipeline B, we either ran the ENCODE pipeline on WT replicates and KO replicates separately (for Pipeline A), or we ran the ENCODE pipeline on all WT and KO replicates simultaneously, setting KO replicates as controls (for Pipeline B). For downstream motif analyses, we used the combined “optimal” peak sets output by IDR.

### Motif analyses with MEME-ChIP

In both pipelines, we employed MEME-ChIP^50,51^ from the MEME Suite^10,11^ for motif analysis. We used MEME-ChIP version 4.12.0, except for CentriMo, which we compiled from version 4.11.2 and modified to output negative sequences. MEME-ChIP performs motif discovery with complementary algorithms MEME^7,8^ and DREME^6^, and motif enrichment with CentriMo^9,44^.

We extended MACS2 narrowPeak regions equidistantly from peak summits to create a uniform set of 500.0000 bp centered peaks^50^. Then, we extracted underlying genomic sequences using BEDTools slop (version 2.23.0)^60^ from a repeat-masked genome. We masked the genome with Tandem Repeats Finder (TRF) (version 4.09)^13^ with options `-h -m -ngs` and parameters `2 7 7 80 10 50 500` for mouse (as done originally by Benson^13^), and options `2 5 5 80 10 30 200` for human (as recommended by Frith et al.^24^).

In Pipeline A, we provided the negative control set in addition to the WT set, running MEME, DREME, and CentriMo in differential mode. In ranking known motifs, we ran CentriMo providing only JASPAR database motifs. Differential CentriMo mode ranks motifs according to Fisher E-values. Since the E-value is the p-value (at most 1) times the number of tests, the E-value cannot exceed the number of motifs in the provided database. Once differential CentriMo reaches the maximum E-value, it starts ranking motifs alphanumerically by motif identifier. Therefore, we do not consider the reported, but relatively meaningless, ranks of motifs with non-significant Fisher E-values. In ranking *de novo* motifs, we ran CentriMo providing only MEME and DREME motifs.

### Pooling *de novo* motifs

Each run of MEME or DREME creates new and globally non-unique identifiers for output motifs. This leads to recurring identifiers that refer to different motifs across multiple runs. To consolidate identifiers across multiple MEME and DREME runs, we modified identifiers to reflect the pipeline from which they originate. We then pooled *de novo* motifs across methods and re-ran the CentriMo step of each pipeline, providing the pooled database, allowing for accurate comparisons.

### Assessing similarity of *de novo* motifs to known motifs

For each experiment, we quantified the similarity of *de novo* motifs to the known JASPAR motif using Tomtom^26^. Tomtom compared the *de novo* motifs to the JASPAR motif database through ungapped alignment across columns^26^. Tomtom generated a list of known motif matches, ranked by increasing Bonferroni-corrected p-values. An exact match between a *de novo* motif and a JASPAR motif would result in the JASPAR motif’s ranking first in this list of matches.

### Comparing input to knockout controls

For experiments with associated input controls, we re-ran our known motif and *de novo* motif analyses swapping out KO datasets for input datasets. We compared peaks between sets using UpSet (version 1.4.0) plots^45^, via Intervene (version 0.6.2)^32^, which calculates genomic region overlaps with BEDTools (version 2.26.0)^60^.

## Availability

PeaKO is available at https://peako.hoffmanlab.org. We additionally include Python source code for peaKO and both pipelines at: https://github.com/hoffmangroup/peako. Persistent availability is ensured by [Zenodo](https://zenodo.org/), in which we have deposited the version of our code we used (https://doi.org/10.5281/zenodo.3338324), its downstream CentriMo and peaKO outputs (https://doi.org/10.5281/zenodo.3338330), and our changes to the CentriMo source code and the Linux x86-64 binary that we used (https://doi.org/10.5281/zenodo.3356995). All source code is licensed under a [GNU General Public License, version 3 (GPLv3)](https://www.gnu.org/licenses/gpl-3.0.en.html), except for CentriMo, which retains its original license.

## Authors’ contributions

Conceptualization, C.V.; Data Curation, D.D. and C.V.; Methodology, D.D., C.V., and M.M.H.; Software, D.D. and C.V.; Visualization, D.D., C.V, and M.M.H.; Writing — Original Draft, D.D.; Writing — Review & Editing, D.D., C.V., and M.M.H.;Resources, M.M.H.;Funding Acquisition, M.M.H.;Supervision, C. V. and M.M.H.

## Acknowledgments

We thank Christopher K. Glass and Verena Link for providing SRF datasets. We thank Carl Virtanen, Zhibin Lu, and Qun Jin (Bioinformatics and High Performance Computing Core, University Health Network).

